# CRISPR Screens Identify Essential Cell Growth Mediators in BRAF-inhibitor Resistant Melanoma

**DOI:** 10.1101/2019.12.16.876631

**Authors:** Ziyi Li, Binbin Wang, Shengqing Gu, Peng Jiang, Avinash Sahu, Chen-Hao Chen, Tong Han, Sailing Shi, Xiaoqing Wang, Nicole Traugh, Hailing Liu, Yin Liu, Qiu Wu, Myles Brown, Tengfei Xiao, Genevieve M. Boland, X. Shirley Liu

## Abstract

BRAF is a serine-threonine kinase that harbors activating mutations in ∼7% of human malignancies and ∼60% of melanomas. Despite initial clinical responses to BRAF inhibitors (BRAFi), patients frequently develop drug resistance. To identify candidate therapeutic targets for BRAFi-resistant melanoma, we conducted CRISPR screens in melanoma cells harboring an activating BRAF mutation that had also acquired resistance to BRAFi. The screens identified pathways and genes critical for BRAFi resistance in melanoma cells. To investigate the mechanisms and pathways enabling resistance to BRAFi in melanomas, we integrated expression data, ATAC-seq, and CRISPR screen results. We identified the JUN family of transcription factors and the ETS family transcription factor ETV5 as key regulators of CDK6 that enabled resistance to BRAFi in melanoma cells. Our findings reveal genes whose loss of function conferred resistance to a selective BRAF inhibitor, providing new insight into signaling pathways that contribute to acquired resistance in melanoma.

## Introduction

Melanoma is an aggressive malignancy with a poor prognosis. The median survival for patients with stage IV melanoma ranges from 8 to 18 months after diagnosis, depending on the substage [1]. Somatic mutations in BRAF, most commonly V600E or V600K [2], are the most frequently identified cancer-causing mutations in melanoma, and recurrently appear in colorectal cancer, non-small cell lung carcinoma, and many other cancers [3]. BRAF encodes a protein belonging to the RAF family of serine/threonine protein kinases. This protein plays a role in regulating the ERK signaling pathway, which affects cell division, differentiation, and cell death [4]. The RAS–RAF–MEK–ERK pathway mediates intracellular responses to growth signals and plays an essential role in tumor progression and metastasis [5].

The frequency of BRAF mutations in metastatic melanoma motivated the development of small molecules targeting mutant BRAF [4]. Early trials indicated that BRAFi treatment showed great promise as a therapeutic strategy for melanomas harboring activating *BRAF* V600E mutations, and was associated with high levels of response [6-8]. BRAF inhibitors vemurafenib and dabrafenib led to improved progression-free survival (PFS) and/or overall survival (OS) versus chemotherapy alone and were approved for the treatment of BRAF-mutant metastatic melanoma [9]. Although a subset of BRAF-mutant cancers respond to small molecule inhibitors of BRAF, the disease usually relapses with acquired resistance [10].

Multiple mechanisms of acquired resistance have been reported. The appearance of BRAF amplifications, BRAF splice variants, and secondary mutations in BRAF such as L514V and L505H can confer resistance to BRAFi [7, 11, 12]. Hyper-activation of components in the RTK-RAS-ERK pathway [13, 14] and persistent expression of the RTK platelet-derived growth factor receptor-β (PDGFRβ) or insulin growth factor-1 receptor (IGF-1R) [13, 15] also led to BRAFi resistance. Activation of other growth pathways, such as mTOR and PI3K, have also been implicated in acquired resistance to BRAFi [16, 17]. The mechanisms of acquired resistance that occur outside of the BRAF gene represent possible targets for combination therapies to counteract BRAFi resistance.

Most tumors, including melanoma, are considered a disease of abnormality in the cell cycle [18]. In melanoma, Cyclin D1 amplification rate is 11%, and this increases to 17% in BRAF V600E melanoma, suggesting a potential role of cyclin D1 in intrinsic resistance to BRAF inhibitors [19]. Increased CDK4 activity also occurs in the majority of melanomas, and CDK4 has been implicated in BRAF inhibitor resistance [19]. Previous studies demonstrated that CDK4/6 inhibitors reduced melanoma cell growth and synergized with BRAF and MEK inhibitors [20-22]. These studies promoted the clinical trials of combined inhibition of BRAF and CDKs. However, it is unknown whether the efficacy of combined pan-CDK4/6 inhibitors with BRAFi is more through CDK4 or CDK6. Studies on the mechanisms of BRAFi resistance will yield important information about the signaling pathways of melanoma pathogenesis as well as how to circumvent this resistance and improve efficacy of drugs.

In order to systematically investigate BRAFi resistance mechanism in melanoma, we conducted a series of experiments in BRAF (V600E) cell lines that had obtained resistance to the BRAFi PLX4032 following chronic exposure [13]. Specifically, our integrative analyses of CRISPR screens, transcriptome and epigenetic profiling, revealed pathways and genes associated with BRAFi resistance and tested candidate combination treatments to counter BRAFi resistance.

## Results

### CRISPR knockout screens in a BRAF-mutant BRAFi-resistant melanoma cell line

To identify genes whose loss of function may counteract resistance to BRAFi, we performed a CRISPR genetic screen in the human melanoma cell line M238R1 [13]. M238R1 is BRAFi-resistant and was derived from long-term high-dose PLX4032 treatment of parental cell line M238 [13]. PLX-4032 and PLX-4720 are both BRAF inhibitors and structurally similar, but PLX-4720 is reported to better inhibit BRAF V600E and to respond better in patient tumor-derived xenografts [23, 24]. To confirm the acquired resistance, we conducted a dose-response assay with PLX-4720 (Figure S1A). The IC50 value of the resistant line was significantly higher than that of the parental line. Previous studies indicated that secondary mutations in BRAF could lead to BRAFi resistance [11]. To rule out the possibility that secondary mutations in BRAF led to BRAFi resistance in M238R1, we sequenced the BRAF coding region. We observed the V600E mutation as expected (Figure S1B), but no other secondary mutations in the BRAF coding region. Meanwhile, there is no BRAF amplification and alternative splicing variants confer BRAFi resistance in this cell lines [25]. This indicates that the drug resistance acquired by M238R1 is not due to a new genetic alteration inside the BRAF coding region.

To identify the genes that confer resistance to BRAF inhibition, we designed a new CRISPR sgRNA library targeting 6000 cancer-related genes (6K-cancer library, TableS 1) based on Cosmic [26] and Oncopanel [27] (Figure 1A and Methods). For each gene, we designed ten 19-bp sgRNAs against the coding region with optimized cutting efficiency and minimized off-target potential using our predictive model [28]. The library contained 1466 sgRNAs against 147 genes essential for cell proliferation as positive controls [29], and 795 non-targeting sgRNAs and 891 sgRNAs targeting AAVS1, ROSA26, and CCR5 as negative controls. We performed two independent, pooled CRISPR screens by transducing a 6K-cancer library of lentivirus to the BRAFi-resistant cells M238R1 (Figure 1B). After viral transduction, we treated the melanoma cells with DMSO or 1uM PLX-4720, an optimal dose chosen based on our preliminary tests (Figure S1A). After 14 days of culturing, we harvested cells from the different treated groups and extracted genomic DNA for PCR the region containing sgRNAs. Then we quantified the abundance of sgRNAs through next-generation sequencing (NGS).

**Figure 1.**
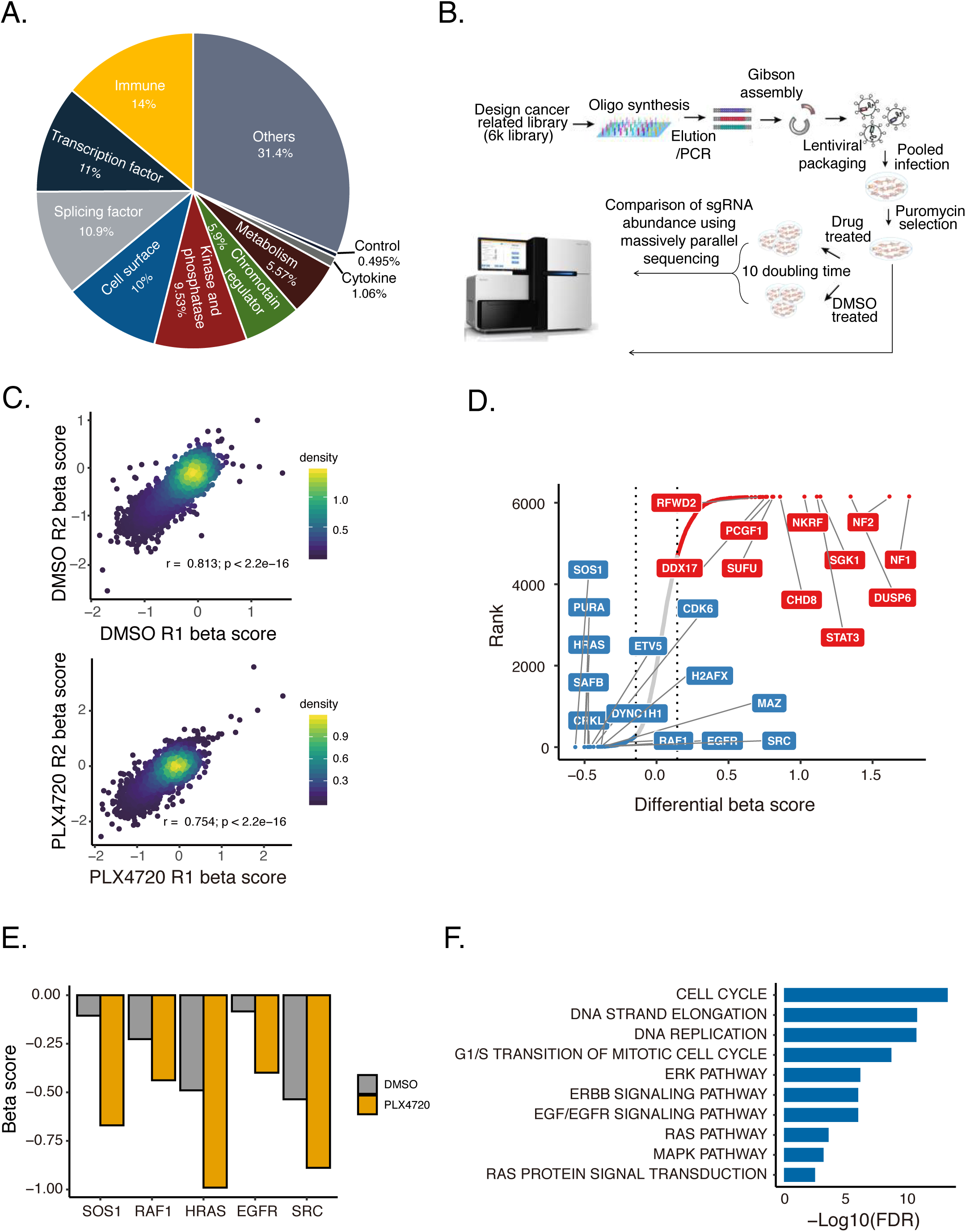
Pooled CRISPR/Cas9-based screens performed in a BRAFi resistant melanoma cell line. **A**. Category of 6K-cancer sgRNA library. **B**. Schematic representation of the workflow for CRISPR screens performed in M238R1 melanoma cells. **C**. Pearson correlation of beta score between two replicates of CRISPR screen data under the treatment of DMSO (top panel) and PLX4720 (bottom panel) in the M238R1 cell line. **D**. Rank of the differential beta score between PLX treatment and vehicle. The two vertical lines indicate +/-1 standard deviation of the difference between treatment and control beta scores. Red dots are genes whose beta score increased after treatment. Blue dots are genes whose beta score decreased after treatment. Gray dots are genes whose beta score did not change significantly between different conditions. **E**. Beta scores of SOS1, RAF1, HRAS, EGFR, and SRC in the PLX4720 condition and DMSO condition. **F**. Pathway enrichment analysis of the essential 322 genes whose β score decreased upon the BRAFi treatment compared to DMSO treatment.

Screen data were analyzed by MAGeCK-VISPR, a statistical algorithm developed for CRISPR screen analyses [30]. MAGeCK-VISPR compares the sgRNA abundance of all of the sgRNAs targeting a gene across different conditions and assigns each gene a log fold-change “beta score (β)” of essentiality in each condition compared with Day 0 control. A positive β-score indicates that silencing corresponding gene provides a growth advantage under the positive selection. In contrast, the negative β-score indicates that silencing the gene confers a growth or survival disadvantage under the negative selection. Replicate screen from the duplicate transductions showed a good correlation at the gene level (Figure 1C). To assess the initial quality of our screen, we check the mapping ratio, the number of missed sgRNAs, and the evenness of sgRNAs (Figure S2). The majority of library was maintained in the viral transduction, with a small amount of missing sgRNA library constructs (Figure S2B). All these results indicated that the screens functioned as designed.

Most genes that were positively or negatively selected behaved similarly in the control and treatment conditions (Table S2). Genes positively selected in both conditions were enriched for known tumor suppressors, such as NF1, NF2 as expected (Figure S3A and B).Consistent with prior work, essential genes highly overlapped between different conditions strongly enriched for roles in fundamental biological processes, such as gene expression, RNA processing, and translation (Figure S3C and D). These results are consistent with a properly functioning CRISPR screen.

### Identification of genes essential specifically for growth of cells resistant to PLX-4720

To explore which genes might play a role in the BRAFi-resistance, we performed further analysis of CRISPR screen data using MAGeCKFlute [29]. MAGeCKFlute facilities comparison of β score between different conditions. We adopted a “quantile matching” approach to robustly estimate σ, which is the standard deviation of the differential β score (Figure S4A). We identified genes whose β score decreased in the presence of BRAFi treatment compared to DMSO treatment (Figure S4B and Table S2). Then, we selected 322 candidates whose disruption does not normally affect survival but becomes lethal in the drug treatment condition. We ranked the identified hits by the change of the β score (Figure 1D). Here, we labeled the top 10 genes, such as *SOS1, PURA, HRAS, SAFB, CRKL, ETV5, <u>CDK6</u>, DYNCH1, H2AFX* and *MAZ*. Among these 322 candidate genes, *HRAS, SRC, SOS1, EGFR*, and *RAF1* were previously reported to be involved in BRAFi resistance [31, 32] (Figure 1E).

To further understand the pathways conferring BRAFi-resistance, we performed GO/GSEA/pathway analyses with the 322 candidate genes (Figure 1F). Among the network of genes whose β score decreased after drug treatment, we found that the ERBB2 signaling pathway, RAS pathway, ERK pathway, MAPK pathway, and EGFR signaling pathway are highly enriched. These results are consistent with previous studies [13, 31, 33, 34]. Besides these known pathways, cell-cycle genes, and G1/ transition S of mitotic cell cycle were the most enriched newly discovered class (Figure 1F), represented by CDK6, CCND1, PSMB1, and RRM2.

### CDK6 confer resistance to BRAF inhibition in melanoma cells

We next sought to determine whether any genes whose upregulation confers resistance to BRAF inhibition in melanoma cells. To assess this, we analyzed previously generated gene expression profiles in parental versus resistant cells treated with PLX4720 or treated with DMSO [13]. In the sensitive cells, PLX4720 induced widespread changes in gene expression (Figure S5A). Our analysis showed that the MAPK signaling pathway and the PI3K-AKT pathway were down-regulated, consistent with previous studies [13, 14] (Figure S5B). The resistant line exhibited fewer differentially expressed genes upon PLX4720 treatment (Figure S5C). We next analyzed the genes that were differentially expressed between the resistant line and the parental line upon PLX4720 treatment. Under BRAFi treatment, there are 1,374 up-regulated and 1,574 down-regulated genes in resistant cells relative to sensitive cells (Figure 2A and Table S3). Our re-analyses confirmed the previously reported overexpression of *KIT, MET, EGFR*, and *PDGFRB* in M238R1 relative to the parental line [13]. In addition, we found that the cell cycle genes *CDK6, CCND1*, and transcription factor (TF) *JUN* were up-regulated in resistant cells compare to the parental cells (Figure 2A).

**Figure 2.**
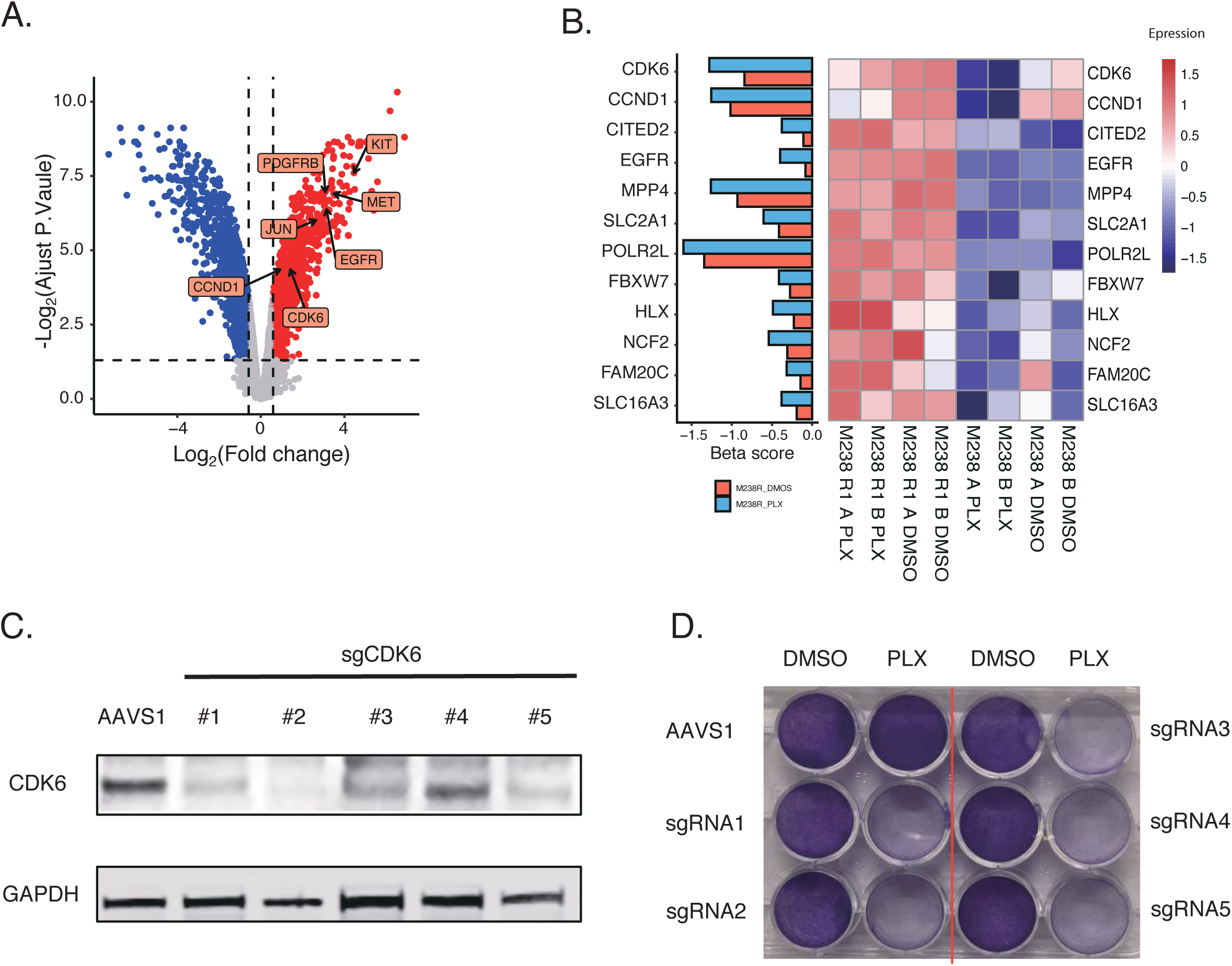
Loss of CDK6 sensitizes cells to BRAFi treatment in M238R1. **A.** Volcano plot showing differentially expressed genes between M238R1 and its parental cell line under the treatment of PLX. The horizontal and vertical lines indicate the cutoff (Fold change >= 1.5; FDR <= 0.05) of differential genes. **B.** Beta score of the screen (left panel) and expression (right panel) of the intersect genes which are more essential in the BRAFi treatment condition and upregulated in the BRAFi-resistant cell line. **C.** Western blots were performed to determine the efficiency of CDK6 sgRNAs. GAPDH was used as a loading control. The M238R1 cells were infected with lentiviruses expressing the indicated sgRNAs at low MOI and selected with puromycin. Cell lysates were blotted with the indicated antibodies. **D.** Loss of CDK6 sensitizes cells to BRAFi treatment in clonogenic assay. Images of colonies in colony formation assay were presented. Results are representative of duplicate biological experiments.

We hypothesized that genes with elevated expression in BRAFi resistant cells, as well as the loss of function restored the drug sensitivity, may be responsible for the resistance phenotype. We next integrated the expression results and CRISPR screen results to identify the dysregulated genes related with BRAFi resistance. Within the 322 genes whose depletion sensitize cells to BRAFi, there are 12 genes, including CDK6, specifically over-expressed in BRAFi-resistant cells (Figure 2B). This suggests that 21 genes might be associated with the resistance to BRAFi and mediate cell proliferation in the resistance line.

To explore the potential druggable targets for the BRAFi-resistant cells, we further filtered the candidate gene with DGIdb [35]. DGIdb is a carefully curated database of published information on drug-gene interactions and the druggable genome. It offers user-friendly functions for browsing, searching, and filtering. DGIdb identified CDK6 as a potential druggable target with the FDA approved drugs for BRAFi-resistant cells. CDK6 is regulated by Cyclin D proteins and Cyclin-dependent kinase inhibitor proteins. Altered expression of these cell cycle genes has been observed in multiple human cancers [36, 37]. CDK6-targeting sgRNAs were markedly depleted in the PLX-4720 condition compared to the DMSO condition (Figure S6A), suggesting that loss-of-function of CDK6 can cause cells sensitive to PLX-4720. To validate this result from the initial screen, we used five independent sgRNAs to knockout CDK6 in the M238R1 cell line (Figure 2C). Consistent with our screen data, CDK6 knockout cells showed increased sensitivity to PLX-4720 in long-term colony-formation viability assays (Figure 2D). Most tumors including melanoma have an abnormal G1-to-S transition, mainly due to dysregulation of CDKs activities [38, 39]. We wondered if the increased essentiality we observed for CDK6 was a general property of CDKs or was specific to CDK6. We specifically evaluated the changes in essentiality of the other CDKs (Figure S6B). Among all CDKs, only CDK6 is more highly expressed in the resistant cell line compared to the sensitive cell line and becomes more essential in the presence of BRAFi.

### Exploring the Mechanism of Gene Regulation in BRAFi Resistance through Chromatin Changes

Epigenetic changes are important features of cancer cells with acquired drug-resistant phenotypes and may be a crucial contributing factor to the development of resistance. To model the epigenetic features associated with BRAFi resistance, we used ATAC-Seq to compare the chromatin accessibility [40] difference between the resistant and parental lines treated with PLX-4720. On average, we sequenced each sample at ∼50 million PE150 fragments and observed ∼89% uniquely mapped ratio (Table S4). We evaluated the quality of deep-sequencing data in diverse sections, such as including the uniquely mapped reads, PCR bottleneck coefficient (PBC) score, High quality peaks number, fraction of non-mitochondrial reads in peak region (FRiP), peaks overlapping with union of DNaseI peaks (DHS) (Figure S7). The ATAC-seq profiles showed the high-quality features according the criteria defined by Cistrome database, which is a data portal for more than 8,000 ChIP-Seq and chromatin accessibility data in human and mouse [41].

In total, 113,725 high-confidence open chromatin regions (or peaks) were identified in the parental line, and 96,038 peaks were identified in resistant line. Of the distinct peaks, we identified the peaks more accessible in parental cells (M238-specific peaks), and the peaks more accessible in resistant cells (M238R1-specific peaks) (Figure 3A and Table S5). Analyzing peaks of accessible chromatin in aggregate provides estimates of the enrichment of transcription factor (TF) binding [42]. M238R1-specific peaks are enriched for genomic locations bound by the AP-1 superfamily, including ATF3, JUNB, AP-1, BATF and JUN (Figure 3B). To investigate the relationship between activated TFs and their target genes, we integrated the ATAC-seq results with gene expression results. We identified the genes that were up-regulated in M238R1 treated with BRAFi and also associated with M238R1-sepecific peaks. These genes are related to EGFR signaling, epithelial cell proliferation, skin development, and angiogenesis (Figure 3C), which are fundamental biological processes of melanoma development. Therefore, analysis of the ATAC-seq data in conjunction with the expression data revealed a set of TFs and their target genes that are associated with BRAFi resistance.

**Figure 3.**
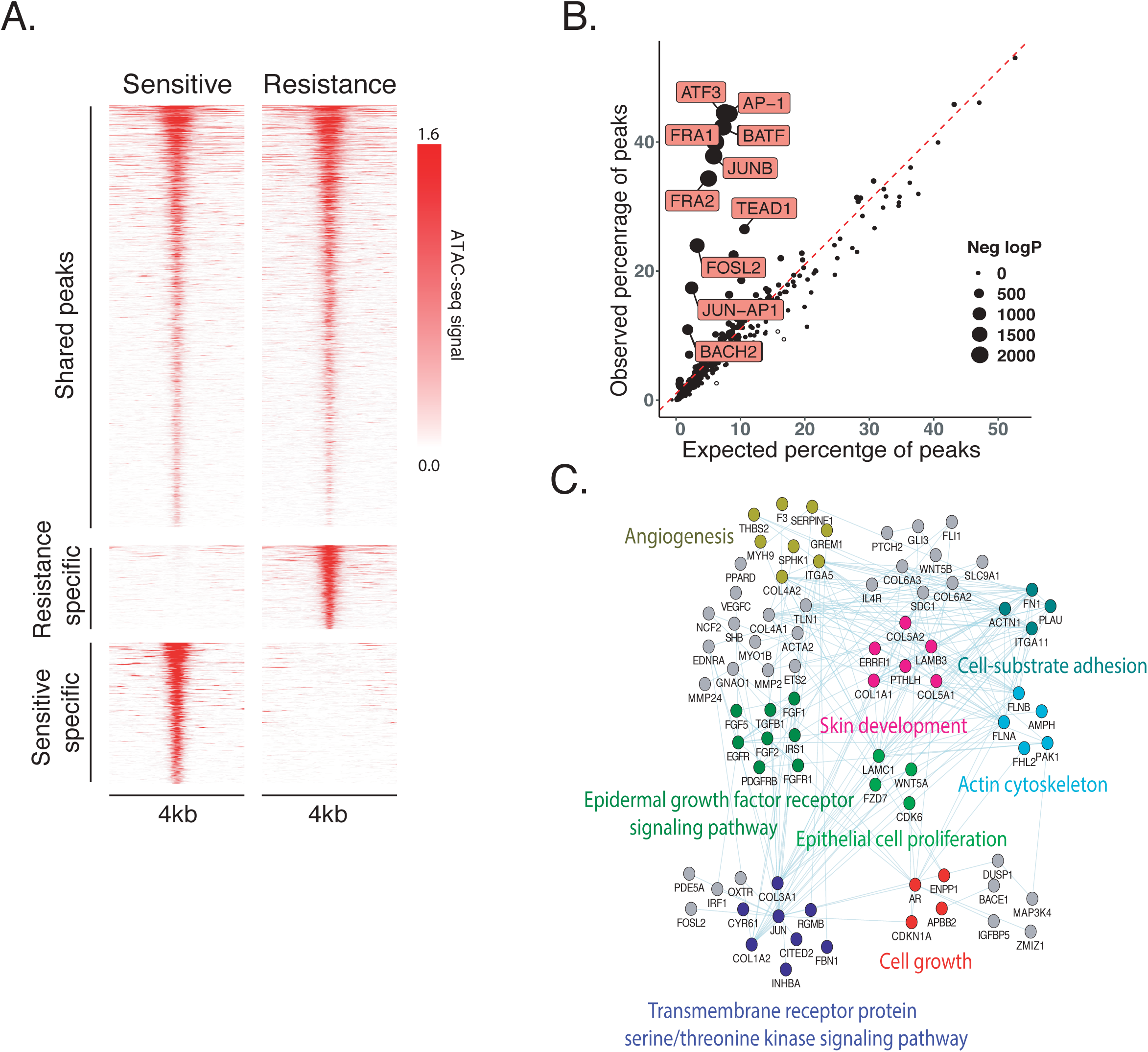
The differences of DNA accessibility between sensitive and resistant cells. **A.** Genome-wide density plots showing that specific and shared ATAC-Seq peaks in BRAFi sensitive and resistant cell lines treated with PLX. Each row represents one peak. The color represents the intensity of chromatin accessibility. Peaks are aligned at the center of regions. **B.** TF motif enrichment. Expected (x axis) versus observed (y axis) percentages of M238R1-specific overlapping each TF binding site annotation. **C.** Network view of the genes which up-regulated in resistant lines treated with BRAFi and also associated with M238R1-sepecific peaks. Here, nodes represent genes and an edge connecting two genes if both are in the same pathway. The pathway information is extracted from the GeneMANIA database [66].

### Identification of the JUN family and ETV5 as key regulators of CDK6

To identify the transcription factors that regulate CDK6 expression, we used the Cistrome ToolKit [41]. The Toolkit allows users to find the factors which might regulate the user-defined genes through public ChIP-seq (protein factors and histone marks), chromatin accessibility (DNase-seq and ATAC-seq) data. We found the AP-1 superfamily JUN, JUNB, and BATF as the putative transcription factors regulating CDK6 (Figure 4A), consistent with previous studies [43, 44]. While all of the transcription factors might regulate CDK6, both expression level (Figure 2A) and chromatin accessibility (Figure 4B) of JUN are higher in the resistant cells. JUN upregulation is a common response to BRAF inhibitor treatment in clinically treated patient tumors and acts as a key mediator of the drug resistance [45, 46]. In addition, JUN is required for cell cycle progression through G1 [47]. As *CDK6* knockout restored sensitivity to BRAFi treatment in M238R1 cells (Figure 2C and D) and *CDK6, JUN* were up-regulated in resistant cells compare to the parental cells, we concluded that dysregulation of CDK6 by JUN mediated resistance to BRAF inhibition in melanoma cells.

**Figure 4.**
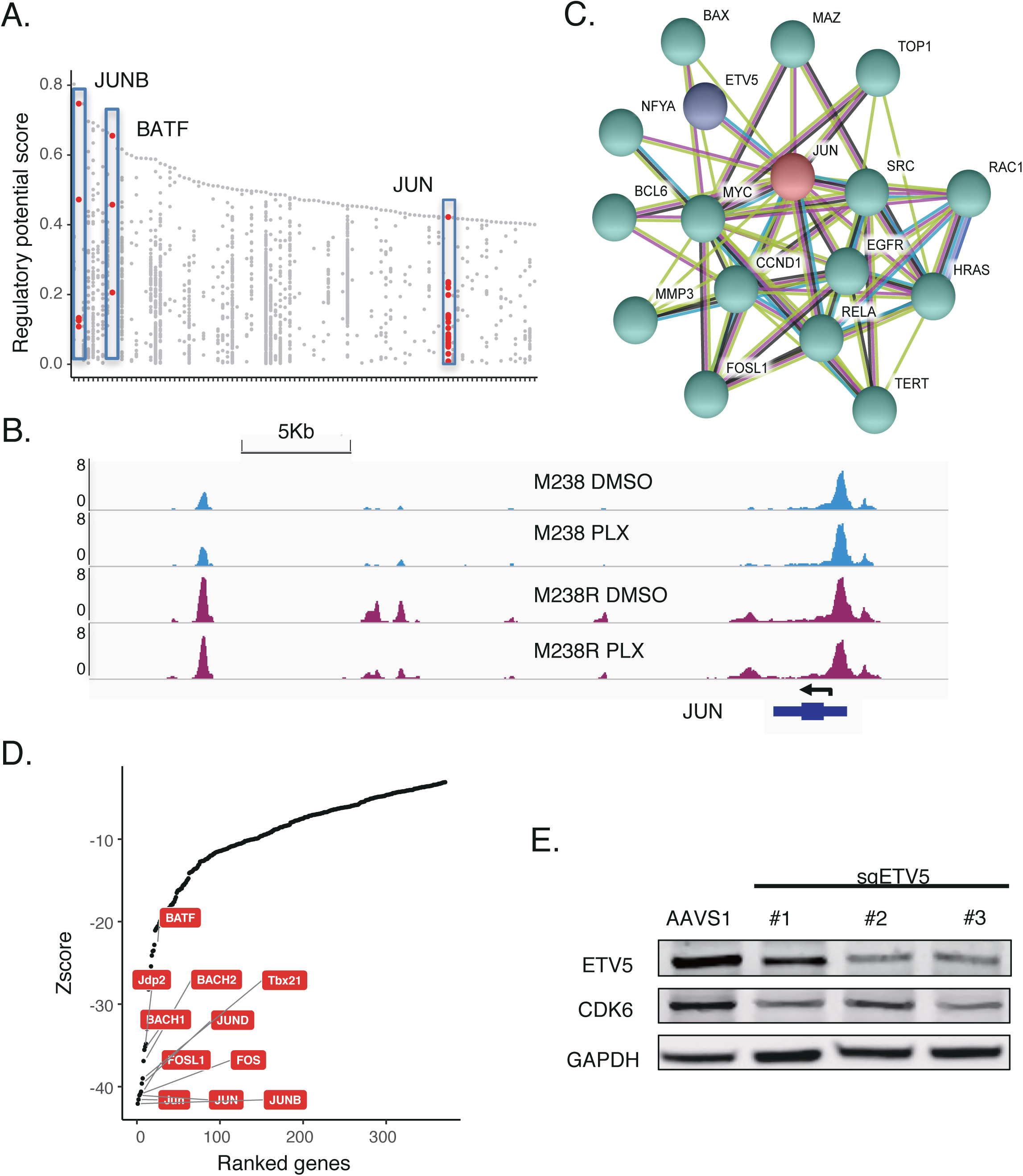
Deficiency of CDK6 or ETV5 combined with PLX4720 inhibit cell proliferation of BRAFi-resistant cells. **A**. Factors which potentially regulate CDK6 are showed in this plot. The y-axis represents the regulatory potential (RP) score which were calculated using Cistrome Data Browser Toolkit. The x-axis represents different factors. Dots in an x-axis line means the same factor. **B**. Browser representation of the region near JUN from ATAC-seq of M238 and M238R1 with different treatment conditions. **C**. Interaction of JUN and genes whose essentiality increased after PLX treatment. Interaction partners of JUN was predicted using STRING database. Colored lines indicate different sources of evidence for each interaction. JUN and ETV5 were individually labeled by the different colors to distinguish them with other genes. **D**. Rank plot of the TF whose motif enriched in the ETV5 Chip-seq peaks. The Zscores are calculated according to their sequence logo similarity using Cistrome MDSeqPos [40]. For the “Zscore” with negative number, the smaller ones mean significantly enriched. **E**. Validation of ETV5 knockout (KO) in M238R1 cells by western blotting using indicated antibodies.

To assess other genes that might act with JUN to regulate CDK6, we examined the set of genes that physically interact with the JUN protein according to the STRING database and genes whose essentiality increased after BRAFi treatment. We identified ETV5 as being in both of these gene sets (Figure 4C). ETV5 is a member of the ETS family of transcription factors which controls cell cycle gene expression and contributes to tumorigenicity [48]. Increased expression of ETV transcription factors modulates the response to MEK inhibition [49]. Motif enrichment analysis of ChIP-seq data can help us identify transcription factors that cooperate with ETV5. According to the Cistrome Data Browser [41], the JUN motif is enriched ETV5 ChIP-seq peaks, suggesting JUN family might be a co-factor of ETV5 (Figure 4D). Consistent with the hypothesis that ETV5, JUN, and JUNB directly regulate CDK6, these TFs have strong binding around the CDK6 gene (Figure S8C). We found that ETV5 deletion reduced sensitivity to BRAF inhibition by PLX-4720 in melanoma cells and ETV5 was the top hit of the genes that were more essential in the BRAFi treatment condition (Figure 1D). Similar to CDK6, the normalized sgRNA read counts of ETV5 continually decrease in the DMSO treatment or PLX-4720 treatment (Figure S8A and B). Finally, we experimentally validated that the depletion of ETV5 decreases the expression of CDK6 (Figure 4E). These observations suggest that CDK6 mediate resistance to BRAF inhibition by the collaborative regulation of TFs JUN and ETV5, which increased expression of CDK6 and promote the cell proliferation.

### Dual inhibition of BRAF and CDK6 in BRAFi-resistant cell lines

Palbociclib (IBRANCE, Pfizer Inc.) is an inhibitor of CDK4 and CDK6 approved by the FDA in many cancer types [50]. CDK inhibitor, and combination of BRAFi or MEKi or a CDK4 inhibitor significantly suppresses growth and enhances apoptosis in melanoma cells [21, 22].However, the efficacy combination therapy of pan-CDK4/6 inhibitors with BRAFi is more through CDK4 or CDK6,whch remains poorly understood. Here, we first examined the changes in essentiality of the other CDKs (Figure S6B). Among all CDKs, only CDK6 is more highly expressed in the resistant cell line compared to the sensitive cell line and becomes more essential in the presence of BRAFi. Further we assessed the synergy between CDK6 and BRAF inhibition on BRAFi resistant cells. To verify the activity of Palbociclib, we showed that 1µM of palbociclib effectively reduced the phosphorylation of CDK6’s substrate RB1 (Figure 5A). We then treated BRAFi resistant cells with palbociclib and/or PLX-4720 and observed that inhibition of CDK6 sensitized cells to PLX-4720 treatment in a clonogenic assay (Figure 5B). This treatment combination is highly synergistic across a broad range of concentrations according to the Bliss independence model, especially in the resistant lines (Figure 5C, 5D and Figure S9). These results support the potential of CDK6 and BRAF dual inhibition as a therapeutic strategy to overcome BRAFi resistance in our resistant model.

**Figure 5.**
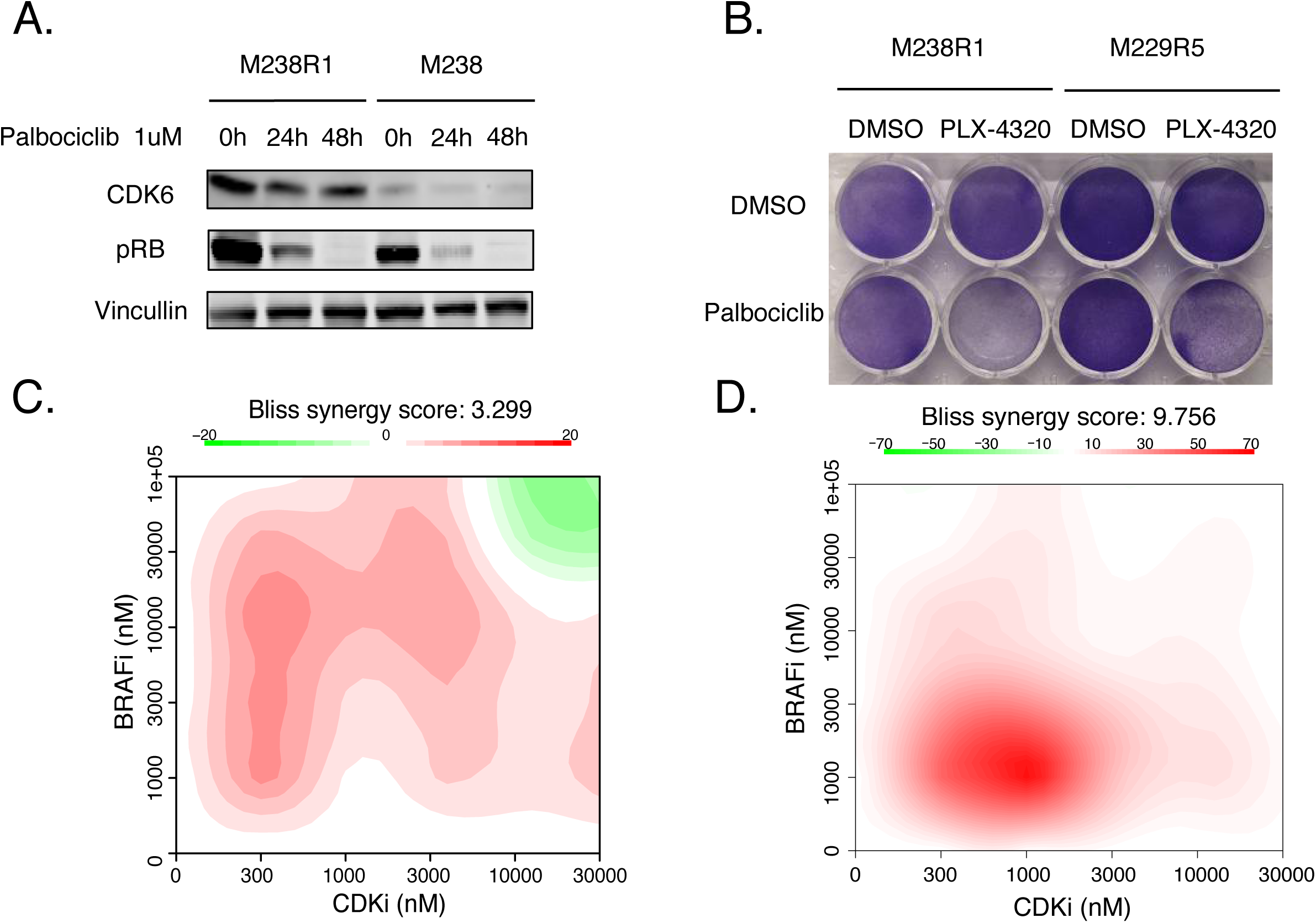
Combination treatment of CDK6i and BRAFi overcame BRAFi resistance in vitro. **A**. Immunoblot of lysates M238 and M238R1 cells that were treated with CDK6 inhibitor at a dosage of 1 uM for 24h and for 72 h. The blot is representative of at least two independent experiments. **B.** The colony formation assay of the combination of palbociclib and PLX-4720 for M238R and M229R5 cell lines. Visualization of the calculated 2D synergy maps of cell line M238R1 (**C**) and M229R5 (**D**). An overall synergy score is calculated as the deviation of phenotypic responses compared to the expected values, averaged over the full dose–response matrix.

### CDK6 expression is negatively associated with overall survival in BRAF-mutant melanomas treated with BRAFi

To determine whether the expression of any validated BRAFi-resistant genes we identified correlated with resistance to BRAF inhibitor therapy in melanomas, we analyzed expression data from two independent cohorts [33, 51]. In cohort one [51], 18 patients were treated either with BRAFi alone (12 patients) or dual BRAFi and MEKi therapies (6 patients). RNA-seq data on serial tumor biopsies of matched pre-treatment and relapsed tumors were available. In cohort two, 22 patients with advanced melanoma were treated with BRAFi (7 patients) or BRAFi plus MEKi (15 patients) [33]. RNA-seq data on pre-treatment, on-treatment, or relapsed tumors were available, although they were not paired. These samples were classified into 3 groups: 14 pre-treatment specimens, 12 on-treatment specimens, and 12 clinical progression specimens. Of the 21 over-expressed genes also identified in our CRISPR screen, *CDK6, CCND1*, and *ETV5* were more highly expressed in the tumors that have relapsed after BRAFi treatment relative to the on-treatment groups (Figure 6A).

**Figure 6.**
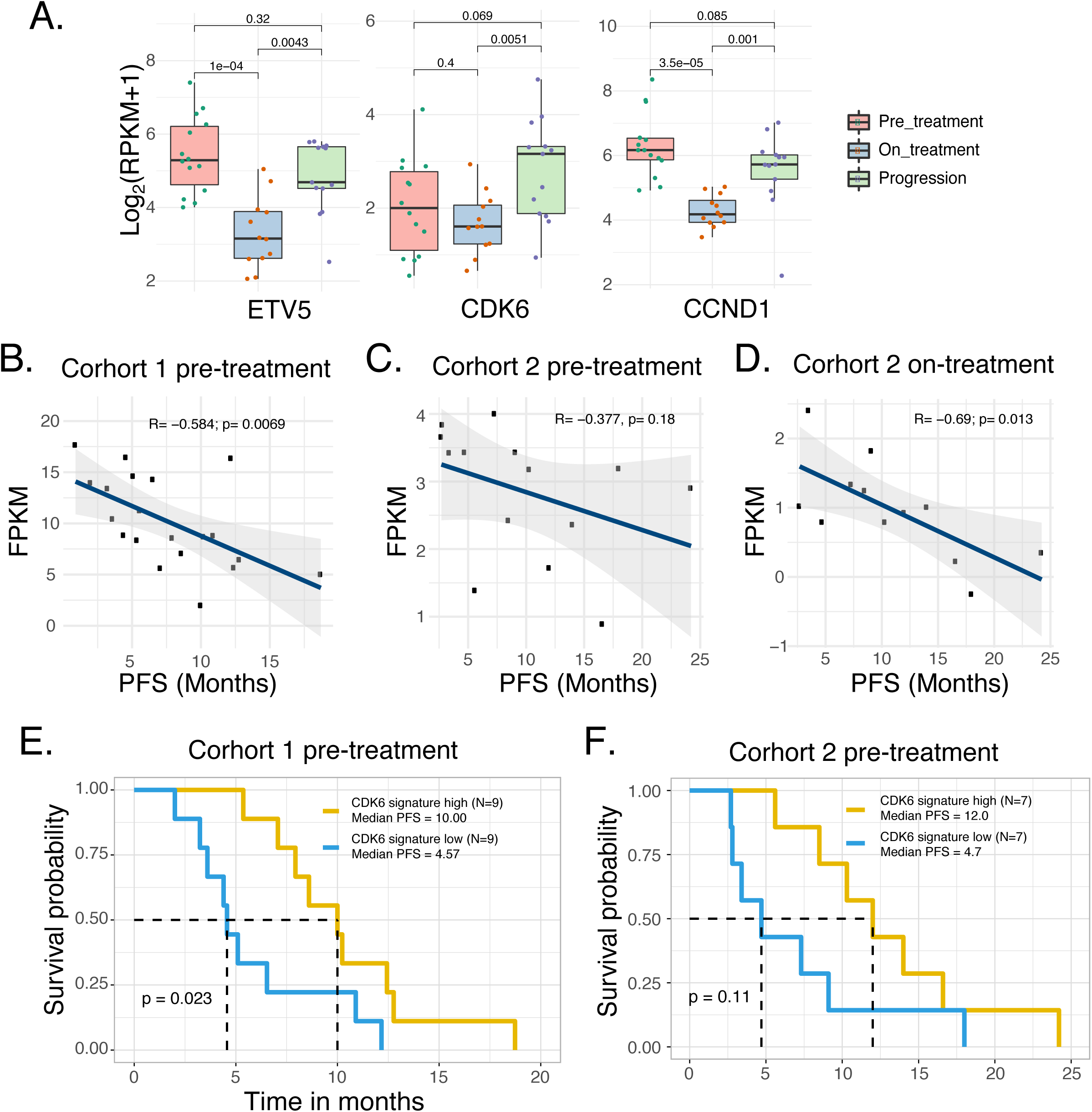
CDK6 and ETV5 expression corelates with cancer progression in patients treated with BARFi. **A**. Expression of ETV5, CDK6, and CCND1 in BRAFi treated patients and progression patients. Correlation of Progression-free survival (PFS) with the CDK6 signature of pre-treatment in cohort 1 (**B**), pre-treatment in cohort 2 (**C**), on-treatment patients in cohort 2(**D**). CDK6 signature overexpression corresponds to worse clinical outcome in a cohort 1 (**E**) and cohort 2 (**F**) patients with melanoma cancer.

We next investigated whether CDK6 upregulation might be associated with clinical resistance in some cases. To facilitate this, we generated a 10-gene CDK6 expression “signature” (Table S6). This 10-gene proliferation signature consists of cell proliferation genes [33] and interaction partners of CDK6 predicted by STRING database. We observed a negative correlation between the CDK6 signature and the progression-free survival (PFS) in samples of both cohorts (Figure 6 B-D). To further clarify the relationship between CDK6 signature and clinical outcome not by the different drug treatment, we separated samples with different drug-treatment condition (BRAFi alone or BRAFi plus MEKi). CDK6 signature was correlated with poor progression-free survival (PFS) of melanoma patients treated with either BRAFi alone or BRAFi plus MEKi (Figure S9 A-C). We used these ten genes to split the samples into CDK6 signature low and CDK6 signature high groups and assessed their prognostic value in melanoma patients of both clinical cohorts. Clinically, melanoma patients classified as CDK6 signature high experienced shorter progression-free survival with respect to CDK6 signature low cases (Figure 6 E and F). Consistent with this, high level of CDK6 signature associated with shorter PFS of the patients either treated with BRAFi alone or BRAFi plus MEKi (Figure S9 D and E). This data suggests that high expression of genes functionally connected to CDK6 associates with poor survival and acquired drug resistance in BRAFi-treated melanoma patients. Overall, these observations provide initial support for the notion that CDK6 upregulation by transcription factors JUN and ETV5 might be associated with clinical resistance to BRAFi in melanoma patients.

## Discussion

Acquired resistance to anticancer agents is frequently encountered in clinical practice. BRAFi-resistance is widely studied but remains a clinical challenge [13, 14, 16, 52].For this reason, it is critical to direct research efforts to investigate the mechanisms underlying drug resistance and design alternative therapeutic strategies to overcome drug resistance. Resistance to kinase inhibitors is often associated with secondary mutations in the target gene, which render the kinase insensitive to the inhibitor [11]. However, in the BRAFi acquired resistant cell line, we did not find secondary mutations in BRAF that could explain the resistance to BRAF inhibitors. Drivers of acquired resistance to BRAF inhibitor therapy are diverse and include mechanisms leading to reactivation of the MAPK pathway [34]. But the M238 R1 was sensitive to PLX4032-induced decreases in the levels of p-MEK1/2 and p-ERK1/2 [13]. Understanding the gene regulation by which cancer cells evade BRAF inhibition may speed the development of new therapeutic strategies in BRAF-mutant melanoma patients and other BRAF-dependent tumors.

Several genome-wide CRISPR pooled screens have uncovered mediators of drug resistance [53, 54]. In this study, we used CRISPR screens to systematically characterize resistance to BRAF inhibitor PLX-4720 in melanoma. Our screen identified both previously known and novel resistance genes to BRAF inhibition. Previously reported genes were identified by our screen, including *CCND1, RAF1, EGFR*, and *SRC* [19, 31, 34]. Among the network of genes whose β score decreased after drug treatment, we also found that the ERBB2 signaling pathway, c-Myc pathway, regulation of RAS family activation, and EGFR signaling pathway represent examples of known pathway-dependent resistance mechanisms [13, 31, 33, 34, 55]. Besides cell-cycle genes were the most enriched newly discovered class (Figure 1F), represented by *<u>CDK6</u>, CCND1, PSMB1*, and *RRM2*. These findings affirm the ability of large-scale functional screens to reveal biologically and clinically relevant drug resistance mechanisms.

Our approach also uncovered depletion of CDK6 and ETV5 restored the sensitivity to BRAF inhibition in BRAFi-resistant cells. To searched for the key regulators of BRAFi resistance, we analyzed gene expression data, chromatin accessibility data, and our CRISPR screen results. Our observations indicate that overexpression of cell cycle gene *CDK6*, which regulated by transcription factors JUN and ETV5, may confer resistance to BRAF inhibition. Indeed, a prior study suggested that overexpression of a single ETS transcription factor conferred resistance to trametinib, suppression of ETV1, ETV4, or ETV5 alone strongly decreased the resistance conferred by CIC deletion [49]. In a previous study, the researchers demonstrated that the inherent resistance to BRAFi/MEKi in melanoma cell lines was associated with a high abundance of JUN [45]. However, JUN family members are not essential for the BRAFi-resistant cell lines. We hypothesize that many JUN family members could collaborate with ETV5 to regulate CDK6, such that the absence of any one member would not lead to cell death. Thus, our integrative analyses of the epigenetic, transcriptional data with genetic screening provided insights into the regulation of BRAFi resistance in melanoma patients.

Palbociclib, an FDA approved drug established to target CDK4/6, has been evaluated in ∼30 different cancer indications [50, 56]. Combining palbociclib with PLX-4120 reduced the proliferation of M238R1 and M229R5, which are BRAFi-resistant melanoma cells. Indeed, prior studies suggested that CDK4/6 inhibition combined with BRAFi inhibited the growth of several melanoma cell lines in vitro and in vivo [20-22]. However, these studies did not determine whether the efficacy of combined CDK4/6 inhibitors with BRAFi was specific to the inhibition of CDK4 or CDK6. Here, we evaluated the essentiality of all CDKs in acquired-BRAFi-resistance cells. Of all the CDKs, only CDK6 is more highly expressed in the resistant cells compared to the sensitive cells, and only CDK6 and becomes more essential in the presence of BRAFi (Figure S5B). Thus, our study demonstrates the feasibility of genome-wide pooled CRISPR-Cas9 knockout screens of resistant cells for uncovering genetic vulnerabilities that may be amenable to therapeutic targeting.

We found that *CDK6* deletion reduced resistance to BRAFi treatment in vitro and demonstrated that the CDK6 inhibitor palbociclib act synergistically with BRAFi to halt cell growth in BRAFi-resistant cell lines. To further demonstrate the potential combination therapy, we tried to generate M238R1 xenografts. However, this effort failed, consistent reports from the lab that derived the resistant cell line (Lo Lab, personal communication). Additional evidence that *CDK6, ETV5* and *JUN* may confer resistance to BRAF inhibition in cancer emerged from our analysis of two independent melanoma cohorts. This analysis revealed high levels of CDK6 and ETV5 in tumors that acquire resistance to BRAFi treatment, thereby providing genetic evidence that these signaling pathways may dysregulate upon BRAF inhibition. A high *CDK6* signature score is associated with the poor progression-free survival of melanoma patients in both clinical cohorts. These observations suggest that elevated global expressions of CDK6, JUN and ETV5 modulate the response to BRAF inhibitor treatment. Our study strengthens this link by demonstrating that a combination of CDK6 inhibitor and BRAF inhibitor can overcome BRAFi resistance.

In conclusion, this study shows that there was a significant increase of CDK6 expression in the BRAFi-resistant cell lines and progressive tumors. Through the loss-of-function screens, epigenetic profiles, and gene expression analysis, we have identified a network that includes CDK6, ETV5, and JUN as the potential mechanism for BRAFi-resistant melanoma cells. Our findings offer new insights into resistance to BRAF inhibitors and support clinical studies of combined BRAF and CDK6 inhibition in a subset of activating BRAF mutations subject to relapse through acquired resistance.

## Materials and methods

### Cell Culture and compounds

Human melanoma paired cell lines were gifts from the Roger Lo lab. Cells were maintained in Dulbecco’s modified Eagle medium (DMEM) with 10% fetal bovine serum, glutamine and 1% penicillin/streptomycin. These BRAFi-sensitive human melanoma cell lines (M series) were established from patient’s biopsies under UCLA IRB approval #02-08-067 [57]. And BRAFi-resistant human melanoma cell lines were derived from long-term high-dose PLX4032 treatment of parental cell line M238 [13]. All cell lines were mycoplasma free. For packaging virus, HEK293T cells were grown in DMEM with 10% FBS, glutamine and 1% penicillin/streptomycin. Stocks of BRAF inhibitor PLX4720 (Catalog No. S1152) and CDK6 inhibitor palbociclib Isethionate (PD0332991, Catalog No. S1579) were purchased from Selleck Chemicals.

### Library design

To design a smaller-scale CRISPR/Cas9 knockout screen library focusing on cancer-related genes, we selected 6000 genes based on reported relevancies with cancers using multiple sources, including Cosmic and Oncopanel (Table S1). For each gene, we designed ten 19nt single-guide RNA (sgRNA) against its coding region with optimized cutting efficiency and minimized off-target potentials. We used sequence features of the spacers to calculate the cutting efficiency score for each sgRNA using our predictive model. We used BOWTIE to map all candidate sgRNAs to hg38 reference genome, and chose those with fewest potential off-targets. We selected the 10 best sgRNAs for each gene based on the considerations above. The library also contains both positive controls and two types of negative controls: non-targeting controls and non-essential-region targeting sgRNAs.

a. Positive controls: we included 1466 sgRNAs targeting 147 positive control genes, which are significantly negatively selected in multiple screen conditions.
b. Non-targeting negative controls: 795 sgRNAs with sequences not found in genome.
c. Non-essential-region-targeting negative controls: 1891 sgRNAs targeting AAVS1, ROSA26, and CCR5, which have been reported as safe-harbor regions where knock-in leads to few detectable phenotypic and genotypic changes.

### Cloning of individual sgRNAs and sgRNA libraries

For the 6K-cancer library, we used the lentiCRISPR v2 vector (also available at Addgene, plasmid #52961) as backbone [58]. We designed ten sgRNAs per gene to target ∼6,000 genes and added non-targeting sgRNAs as controls (Table S1). For library construction, we used a previously published protocol [54]. For individual sgRNA cloning, pairs of oligonucleotides (IDT) with BsmBI-compatible overhangs were separately annealed and cloned into the lentiCRISPR v2 vector using standard protocols [58]. The sequences of individual sgRNAs for *CDK6* and *ETV5* are shown in Table S7.

### Virus production and infection

Lentivirus was generated in HEK293T cells by transfecting cells with packaging DNA plus lenti-CRISPR vectors. For each library to be transfected, we plated HEK293T cells in 25ml of media in a 15 cm tissue culture plate. Typically, 20 μg vector DNA, 15 ug psPAX2 packaging plasmid, 6 ug pMD2.G envelope plasmid and 200 ul transfection reagent X-tremeGENE were used; DNA and transfection reagent X-tremeGENE were pre-diluted in 3 ml serum-free OPTI-MEM individually and then mixed. After 15 min of incubation, the DNA and transfection reagent mixtures were added to HEK293T cells seeded in the dish. After 8-12 h, the media was changed to 25 ml DMEM + 10% FBS+ 1%BSA. Viral supernatant was collected two and three days after transfection, filtered through 0.45-μm membranes, and added to target cells in the presence of polybrene (8 μg/ml, Millipore). After 48h, puromycin (2 μg/ml) was used to treat cells for two days for selection, which eliminated all cells in an uninfected control group.

### Pooled CRISPR screen

For the pooled CRISPR screen, a total of 1.2×10^8^ cells were infected with the pooled lentiviral library at a MOI of 0.3. After puromycin selection, the surviving cells were divided into three groups (day0 control, vehicle, and drug treatment). For the drug treatment group, the cells were treated with 1uM PLX4720. The cells were cultured in medium for ten doubling times and split every 2-3 days before genomic DNA extraction and library amplification.

### Amplification and sequencing of sgRNAs from cells

After cell harvest, DNA was purified using QIAGEN DNeasy Blood & Tissue Kit according to the manufacturer’s instruction. PCR was performed as previously described [58], and the PCR products were sequenced on a HiSeq 2500. Each library was sequenced at 30∼40 million reads to achieve ∼300X average coverage over the CRISPR library. The day 0 sample library of each screen could serve as controls to identify positively or negatively selected genes or pathways.

### CRISPR screen analysis

The CRISPR/Cas9 screening data were analyzed using MAGeCK and MAGeCK-VISPR algorithms [30]. MAGeCK-VISPR uses a metric, “β score”, to measure gene selections. The definition of the β score is similar to the term of ‘log Fold Change’ in differential expression analysis, and β>0 (or <0) means the corresponding gene is positively (or negatively) selected, respectively. We considered a β score of >0.5 or <-0.5 as significant. MAGeCK-VISPR models the gRNA read counts as a negative binomial variable, whose mean value is determined by the sequencing depth of the sample, the efficiency of the gRNA, and a linear combination of β scores of the genes. MAGeCK-VISPR then builds a maximum likelihood (MLE) model to model all gRNA read counts of all samples, and iteratively estimate the gRNA efficiency and gene β scores using the Expectation-Maximization algorithm. Comparison between the drug treatment condition and control condition was performed using MAGeCKFlute [29], which was designed to perform quality control, normalization, gene hit identification and downstream functional enrichment analysis for CRISPR screens.

### Microarray data analysis

The expression profile GSE9340, which was downloaded from Gene Expression Omnibus database, included two BRAFi resistant cell lines (M238R and M229R) and their parental cell lines (M238 and M229). Differential expression analysis was performed using the R package limma [59]. Genes with an absolute fold change >1.5 and false discovery rate (FDR)-adjusted P□<□0.05 were considered significant.

### ATAC-seq

ATAC-seq libraries were prepared according to the previously described Omni-ATAC protocol [60]. After the cells counting, 50,000 cells were resuspended in 1 ml of cold ATAC-seq resuspension buffer (RSB; 10 mM Tris-HCl pH 7.4, 10 mM NaCl, and 3 mM MgCl2 in water). Cells were pelleted by centrifugation at 500 r.c.f at 4 °C for 5 min in a pre-chilled (4 °C) centrifuge. After centrifugation, supernatant was carefully aspirated to leave the cell pellet undisturbed. Cell pellets were then resuspended in 50 μl of ATAC-seq resuspension buffer containing 0.1% NP40, 0.1% Tween-20, and 0.01% digitonin by pipetting up and down three times. This cell lysis reaction was incubated on ice for 3 min. After lysis, 1 ml of ATAC-seq RSB containing 0.1% Tween-20 (without NP40 or digitonin) was added, and the tubes were inverted 3 times to mix. Nuclei were then centrifuged for 10 min at 500 r.c.f. 4 °C centrifuge. Nuclei were resuspended in 50 μl of transposition mix (25 μl 2× TD buffer, 2.5 μl transposase (100 nM final), 16.5 μl PBS, 0.5 μl 1% digitonin, 0.5 μl 10% Tween-20, and 5 μl nuclease-free water) by pipetting up and down six times. Transposition reactions were incubated at 37 °C for 30 min in an Eppendorf ThermoMixer with shaking at 1,000 r.p.m. Tagmented DNA was purified using the MinElute Reaction Cleanup Kit (Qiagen, 28204). The ATAC-seq library preparation was performed as described previously [40]. Then, the concentration of the library was determined using Qubit 3.0 (Life Technologies) and the size distribution was assessed using Agilent 4200 TapeStation system. Libraries were paired-end sequenced (35bp) on an Illumina NextSeq 500.

### ATAC-seq data analysis

Quality control, reads alignment, peak calling were performed by ChiLin [61]. The M238 and M238R peaks were further merged (using the BEDtools [62] ‘merge’ function). BEDtools ‘coverage’ was used to create an input matrix used for detecting differentially accessible, peaks. We assessed the significant change of chromatin accessibility between different groups using the DESeq2 R package [63]. The total count of qualified fragments in each sample was used as the library size. It was defined as significantly changed if the peak showed log2 fold change > 1 and adjust P-value < 0.05. The HOMER tool suite was used for TF motif discovery, by analyzing differential motif enrichment in M238R specific element datasets against all elements (peaks) background. Regulatory potential (RP) scores derived with the BETA algorithm are used to estimate how likely a factor regulates genes [64].

### ChIP-seq data mining in Cistrome Data Browser

We used the Cistrome Data Browser Toolkit function to investigate the transcriptional factors which could regulate CDK6 [41]. This function would return a list of the transcription factors that are most likely to regulate of CDK6. To identify the potential cooperative factors of ETV5, we used the analysis results from the Cistrome Data Browser [41]. ETV5 ChIP-seq data with the high-quality (Cistrome Data Browser ID: 42714) were used to explore the potential cooperative factors of ETV5. In the “QC Motifs” panel, it shows the significantly enriched motifs of other factors in the ETV5 ChIP-seq peaks.

### Western Blot analysis

For western blotting, cells were lysed in RIPA buffer (Santa Cruz Biotechnology) supplemented with phosphatase and protease inhibitor cocktail. Protein concentrations were measured with Thermo Fisher Scientific Bradford Assay (# PI23236). ETV5 Antibody (catalog: ab102010) was purchased from Abcam, and CDK6 Antibody (catalog: sc-7961) was purchased from Santa Cruz Biotechnology. ERK2 Antibody (Santa Cruz Biotechnology, sc-1647) GAPDH (Sigma, G9545), and VINCULIN (Santa Cruz Biotechnology, sc-73614) were used as a loading control. Goat anti-rabbit and Goat anti-mouse secondary antibodies were obtained from LI-COR Biosciences. The fluorescent signals were developed with Odyssey CLX Imaging System (LI-COR Biosciences).

### Cell proliferation and colony formation assays

Response to a single agent-or combination-treatment was assessed by either the CellTiter 96 cell proliferation assay from Promega. Cells were seeded in 96-well plates (2,000 cells per well), and cultured 18 to 24 hours before compound addition. The cells were treated with various concentrations of BRAFi or/and CDK6i for 72 hr and then incubated with CellTiter 96 AQueous One Solution Reagent for 1-4 hr per manufacturer’s protocol before recording the absorbance at 490 nm on SpectraMax M2 (Molecular Devices). All experiments were performed in triplicate. For colony formation assays, cells were seeded in a 24-well plate at a density of 300, allowed to attach for 24 hours at 37°C, and then treated with PLX4720. The cells were maintained at 37°C for two weeks. Colonies of cells were then fixed with cold methanol for 25 minutes and stained with 1% crystal violet.

### Drug synergy analysis

Drug synergy was calculated based on the Bliss independence model using the SynergyFinder R package [65].The Synergy score based on Bliss model.

## Author contributions

SL conceptualized the study, supervised the experiments and data analysis. ZL and BW conceived and designed the study. ZL performed all experiments including the screening, in vitro experiments. SG supervised experiments and provided technical support. BW performed computational analysis of the data. GB. and AS provides the data of the patients in cohort two. CHC designed the CRISPR screen library. TX constructed the CRISPR-sgRNA library. PJ, TH, QW, and SS participated in experiments. PJ, HL, YL, and MB contributed to the discussion. SL, ZL, and BW wrote this manuscript with feedback from all other authors. All authors read and approved the final manuscript.

## Acknowledgments

We thank the Roger Lo Lab for sharing the melanoma cell lines including the parental and resistant lines, and for helpful discussions. This project was supported by grants from the National Natural Science Foundation of China (Grant No. 81872290, 31801185).

## Supplementary material

**Figure S1. BRAF V600E mutation and BRAF inhibitor PLX-4720 resistance**

**A**. Growth curves for parental melanoma cell lines and their isogenic BRAFi-resistant sub-lines. Cells were treated with the PLX4720 for 72 h. **B**. Codons encoding glutamic acid at amino acid position 600 highlighted in red.

**Figure S2. The quality control measurements of the CRISPR screens**

**A.** Read counts and mapping ratio. **B.** Number of missed sgRNAs. **C.** Gini index, which measures read depth evenness within samples. **D.** Violin plot of beta score M238R cells under DMSO and PLX4720 treatment respectively.

**Figure S3. Analysis of positively and negatively selection genes in CRISPR screen** Positively and negatively selected genes in M238R cell line under the DMSO treatment **(A)** and PLX4720 treatment **(B)**. The pathway enrichment analysis of the negatively selected gene in M238R1 cell line treated with DMSO **(C)** and PLX **(D)**.

**Figure S4. Comparison of the genes’ beta score in different conditions**

**A.** Density plot of differential beta scores compared PLX4720 treatment condition with DMSO treatment condition. Delta is used to measure the change of beta score in the two conditions. Delta was calculated by the formula shown in the right panel. If the genes’ differential beta scores are bigger than delta (the red line), theses genes’ essentiality decreased after PLX4720 treatment. Delta is 0.134 in our screen data. Genes’ differential beta scores are smaller than minus delta (the blue line), which indicate theses genes’ essentiality increased after PLX4720 treatment. **B.** The beta score of M238R1 cell line with the treatment of DMSO and PLX4720. The two diagonal lines indicate +/-1 delta of the difference between treatment and control beta scores. Red dots are genes whose beta score increased after treatment. Blue dots are genes whose beta score decreased after treatment.

**Figure S5. The differences of expression between sensitive and resistant lines with or without drug treatment**

**A.** Volcano plot shows the differential expressed genes between the treatment of PLX4720 and DMSO in BRAFi sensitive cell line (M238). **B.** Enrichment results of the down-regulated genes after PLX4720 treated compared with vehicle in M238 parental cell line. **C.** Volcano plot shows the differential expressed genes between the treatment of PLX4720 and DMSO in BRAFi resistant cell line (M238R1).

**Figure S6. The dependency of CDK6 and cell cycle gene in BRAFi-resistant cells between different conditions**

**A**. Boxplot of normalized read count of sgRNAs that target CDK6 in BRAFi resistant M238 cell line in Day0, DMSO and PLX4720 conditions. **P < 0.01, *P < 0.05, two-sided Wilcoxon signed rank test. NS, not significant. **B.** Beta score of CDK genes in different treatment conditions. **C.** Volcano plot shows differential expressed CDK genes between the BRAFi resistant cell line (M238R1) and sensitive cell line with the treatment of PLX4720.

**Figure S7. ATAC-seq data quality control**

**A.** The median sequence quality score. **B**. Uniquely mapped reads are the number of reads with mapping quality above 1. The uniquely mapped ratio is the uniquely mapped reads divided by the total reads. **C.** PCR bottleneck coefficient (PBC) is the locations with only one read divided by unique locations. **D.** The 10-fold confident peaks are the number of peaks called by MACS2, where the fold change is 10. **E.** The fraction of non-mitochondrial reads in peak region (FRiP) score assesses the ChIP-seq signal to noise ratio, which the definition is the fraction of mapped or usable reads that locate in the called peaks. **F.** The DHS ratio of reads is the estimated ratio of reads falling in DNaseI Hypersensitive regions. The red lines indicate the cutoff of good quality data, which was learned from the mass of epigenetic data by the Cistrome Data Browser.

**Figure S8. The dependency and ChIP-seq profiling of ETV5**

**A.** Chip-seq pilled reads of JUN, JUNB, ETV5. Boxplot **(B)** and segment plot **(C)** of normalized read count of sgRNAs that target ETV5 in BRAFi resistant M238 cell line in Day0, DMSO and PLX4720 conditions. **P < 0.01, *P < 0.05, two□sided Wilcoxon signed rank test. NS, not significant.

**Figure S9. Combination synergy assay in vitro**

Dose response of PLX4720 with increasing amounts of Palbociclib for M238R1 **(A)** and M229R5 **(B)** cell lines. An overall synergy score is calculated as the deviation of phenotypic responses compared to the expected values, averaged over the full dose–response matrix. Visualization of the calculated 3D synergy maps of M238R1 **(C)** and M229R5 **(D)** cell lines.

**Supplementary table 1. sgRNA sequences of 6K CRISPR screen library.**

**Supplementary table 2. Beta score of M238R1 CRISPR Screens.**

**Supplementary table 3. Significantly differentially expressed genes in M238R1 treated with PLX-4720.**

**Supplementary table 4. Mapping ratio of ATAC-seq of M238R1 and M238 cell lines.**

**Supplementary table 5. M238R1 specific peaks**

**Supplementary table 6. 10 genes of CDK6 expression “signature”.**

**Supplementary table 7. sgRNA sequences of CDK6 and ETV5.**

## References

[1] Balch CM, Gershenwald Je Fau - Soong S-J, Soong Sj Fau - Thompson JF, Thompson Jf Fau - Atkins MB, Atkins Mb Fau - Byrd DR, Byrd Dr Fau - Buzaid AC, et al. Final version of 2009 AJCC melanoma staging and classification.

[2] Gorden A, Osman I, Gai WM, He D, Huang WQ, Davidson A, et al. Analysis of BRAF and N-RAS mutations in metastatic melanoma tissues. Cancer Research 2003;63:3955–7.

[3] Davies H, Bignell GR, Cox C, Stephens P, Edkins S, Clegg S, et al. Mutations of the BRAF gene in human cancer. Nature 2002;417:949–54.

[4] Peyssonnaux C, Eychene A. The Raf/MEK/ERK pathway: new concepts of activation. Biol Cell 2001;93:53–62.

[5] McCubrey JA, Steelman LS, Chappell WH, Abrams SL, Wong EW, Chang F, et al. Roles of the Raf/MEK/ERK pathway in cell growth, malignant transformation and drug resistance. Biochim Biophys Acta 2007;1773:1263–84.

[6] Fedorenko IV, Paraiso KH, Smalley KS. Acquired and intrinsic BRAF inhibitor resistance in BRAF V600E mutant melanoma. Biochem Pharmacol 2011;82:201–9.

[7] Poulikakos PI, Persaud Y, Janakiraman M, Kong XJ, Ng C, Moriceau G, et al. RAF inhibitor resistance is mediated by dimerization of aberrantly spliced BRAF(V600E). Nature 2011;480:387–U144.

[8] Villanueva J, Infante JR, Krepler C, Reyes-Uribe P, Samanta M, Chen HY, et al. Concurrent MEK2 mutation and BRAF amplification confer resistance to BRAF and MEK inhibitors in melanoma. Cell Rep 2013;4:1090–9.

[9] Larkin J, Del Vecchio M, Ascierto PA, Krajsova I, Schachter J, Neyns B, et al. Vemurafenib in patients with BRAF(V600) mutated metastatic melanoma: an open-label, multicentre, safety study. Lancet Oncol 2014;15:436–44.

[10] Moriceau G, Hugo W, Hong A, Shi H, Kong X, Yu CC, et al. Tunable-combinatorial mechanisms of acquired resistance limit the efficacy of BRAF/MEK cotargeting but result in melanoma drug addiction. Cancer Cell 2015;27:240–56.

[11] Wang J, Yao Z, Jonsson P, Allen AN, Qin ACR, Uddin S, et al. A Secondary Mutation in BRAF Confers Resistance to RAF Inhibition in a BRAF(V600E)-Mutant Brain Tumor. Cancer Discov 2018;8:1130–41.

[12] Villanueva J, Infante JR, Krepler C, Reyes-Uribe P, Samanta M, Chen HY, et al. Concurrent MEK2 Mutation and BRAF Amplification Confer Resistance to BRAF and MEK Inhibitors in Melanoma. Cell Reports 2013;4:1090–9.

[13] Nazarian R, Shi H, Wang Q, Kong X, Koya RC, Lee H, et al. Melanomas acquire resistance to B-RAF(V600E) inhibition by RTK or N-RAS upregulation. Nature 2010;468:973–7.

[14] Wagle N, Van Allen EM, Treacy DJ, Frederick DT, Cooper ZA, Taylor-Weiner A, et al. MAP kinase pathway alterations in BRAF-mutant melanoma patients with acquired resistance to combined RAF/MEK inhibition. Cancer Discov 2014;4:61–8.

[15] Villanueva J, Vultur A, Lee JT, Somasundaram R, Fukunaga-Kalabis M, Cipolla AK, et al. Acquired resistance to BRAF inhibitors mediated by a RAF kinase switch in melanoma can be overcome by cotargeting MEK and IGF-1R/PI3K. Cancer Cell 2010;18:683–95.

[16] Greger JG, Eastman SD, Zhang V, Bleam MR, Hughes AM, Smitheman KN, et al. Combinations of BRAF, MEK, and PI3K/mTOR inhibitors overcome acquired resistance to the BRAF inhibitor GSK2118436 dabrafenib, mediated by NRAS or MEK mutations. Mol Cancer Ther 2012;11:909–20.

[17] Paraiso KHT, Xiang Y, Rebecca VW, Abel EV, Chen YA, Munko AC, et al. PTEN Loss Confers BRAF Inhibitor Resistance to Melanoma Cells through the Suppression of BIM Expression. Cancer Research 2011;71:2750–60.

[18] Hanahan D, Weinberg RA. The hallmarks of cancer. Cell 2000;100:57–70.

[19] Smalley KSM, Lioni M, Palma MD, Xiao M, Desai B, Egyhazi S, et al. Increased cyclin D1 expression can mediate BRAF inhibitor resistance in BRAF V600E-mutated melanomas. Molecular Cancer Therapeutics 2008;7:2876–83.

[20] Martin CA, Cullinane C, Kirby L, Abuhammad S, Lelliott EJ, Waldeck K, et al. Palbociclib synergizes with BRAF and MEK inhibitors in treatment naive melanoma but not after the development of BRAF inhibitor resistance. Int J Cancer 2018;142:2139–52.

[21] Yoshida A, Lee EK, Diehl JA. Induction of Therapeutic Senescence in Vemurafenib-Resistant Melanoma by Extended Inhibition of CDK4/6. Cancer Res 2016;76:2990–3002.

[22] Yadav V, Burke TF, Huber L, Van Horn RD, Zhang Y, Buchanan SG, et al. The CDK4/6 inhibitor LY2835219 overcomes vemurafenib resistance resulting from MAPK reactivation and cyclin D1 upregulation. Mol Cancer Ther 2014;13:2253–63.

[23] Tsai J, Lee JT, Wang W, Zhang J, Cho H, Mamo S, et al. Discovery of a selective inhibitor of oncogenic B-Raf kinase with potent antimelanoma activity. Proc Natl Acad Sci U S A 2008;105:3041–6.

[24] Bollag G, Hirth P, Tsai J, Zhang J, Ibrahim PN, Cho H, et al. Clinical efficacy of a RAF inhibitor needs broad target blockade in BRAF-mutant melanoma. Nature 2010;467:596–9.

[25] Shi HB, Moriceau G, Kong XJ, Lee MK, Lee H, Koya RC, et al. Melanoma whole-exome sequencing identifies B-V600E-RAF amplification-mediated acquired B-RAF inhibitor resistance. Nature Communications 2012;3.

[26] Forbes SA, Beare D, Boutselakis H, Bamford S, Bindal N, Tate J, et al. COSMIC: somatic cancer genetics at high-resolution. Nucleic Acids Res 2017;45:D777–D83.

[27] Garcia EP, Minkovsky A, Jia YH, Ducar MD, Shivdasani P, Gong X, et al. Validation of OncoPanel A Targeted Next-Generation Sequencing Assay for the Detection of Somatic Variants in Cancer. Archives of Pathology & Laboratory Medicine 2017;141:751–8.

[28] Xu H, Xiao T, Chen CH, Li W, Meyer CA, Wu Q, et al. Sequence determinants of improved CRISPR sgRNA design. Genome Res 2015;25:1147–57.

[29] Wang B, Wang M, Zhang W, Xiao T, Chen CH, Wu A, et al. Integrative analysis of pooled CRISPR genetic screens using MAGeCKFlute. Nat Protoc 2019.

[30] Li W, Koster J, Xu H, Chen CH, Xiao T, Liu JS, et al. Quality control, modeling, and visualization of CRISPR screens with MAGeCK-VISPR. Genome Biol 2015;16:281.

[31] Girotti MR, Pedersen M, Sanchez-Laorden B, Viros A, Turajlic S, Niculescu-Duvaz D, et al. Inhibiting EGF receptor or SRC family kinase signaling overcomes BRAF inhibitor resistance in melanoma. Cancer Discov 2013;3:158–67.

[32] Antony R, Emery CM, Sawyer AM, Garraway LA. C-RAF Mutations Confer Resistance to RAF Inhibitors. Cancer Research 2013;73:4840–51.

[33] Kwong LN, Boland GM, Frederick DT, Helms TL, Akid AT, Miller JP, et al. Co-clinical assessment identifies patterns of BRAF inhibitor resistance in melanoma. J Clin Invest 2015;125:1459–70.

[34] Corcoran RB, Ebi H, Turke AB, Coffee EM, Nishino M, Cogdill AP, et al. EGFR-Mediated Reactivation of MAPK Signaling Contributes to Insensitivity of BRAF-Mutant Colorectal Cancers to RAF Inhibition with Vemurafenib. Cancer Discovery 2012;2:227–35.

[35] Cotto KC, Wagner AH, Feng YY, Kiwala S, Coffman AC, Spies G, et al. DGIdb 3.0: a redesign and expansion of the drug-gene interaction database. Nucleic Acids Res 2018;46:D1068–D73.

[36] Sherr CJ, McCormick F. The RB and p53 pathways in cancer. Cancer Cell 2002;2:103–12.

[37] Malumbres M, Barbacid M. Cell cycle, CDKs and cancer: a changing paradigm. Nat Rev Cancer 2009;9:153–66.

[38] Malumbres M, Barbacid M. Is Cyclin D1-CDK4 kinase a bona fide cancer target? Cancer Cell 2006;9:2–4.

[39] Malumbres M, Barbacid M. To cycle or not to cycle: A critical decision in cancer. Nature Reviews Cancer 2001;1:222–31.

[40] Buenrostro JD, Giresi PG, Zaba LC, Chang HY, Greenleaf WJ. Transposition of native chromatin for fast and sensitive epigenomic profiling of open chromatin, DNA-binding proteins and nucleosome position. Nat Methods 2013;10:1213–8.

[41] Zheng R, Wan C, Mei S, Qin Q, Wu Q, Sun H, et al. Cistrome Data Browser: expanded datasets and new tools for gene regulatory analysis. Nucleic Acids Res 2019;47:D729–D35.

[42] Corces MR, Granja JM, Shams S, Louie BH, Seoane JA, Zhou W, et al. The chromatin accessibility landscape of primary human cancers. Science 2018;362.

[43] Kollmann K, Heller G, Sexl V. c-JUN prevents methylation of p16(INK4a) (and Cdk6): the villain turned bodyguard. Oncotarget 2011;2:422–7.

[44] Schreiber M, Kolbus A, Piu F, Szabowski A, Mohle-Steinlein U, Tian JM, et al. Control of cell cycle progression by c-Jun is p53 dependent. Genes & Development 1999;13:607–19.

[45] Ramsdale R, Jorissen RN, Li FZ, Al-Obaidi S, Ward T, Sheppard KE, et al. The transcription cofactor c-JUN mediates phenotype switching and BRAF inhibitor resistance in melanoma. Science Signaling 2015;8.

[46] Titz B, Lomova A, Le A, Hugo W, Kong XJ, ten Hoeve J, et al. JUN dependency in distinct early and late BRAF inhibition adaptation states of melanoma. Cell Discovery 2016;2.

[47] Wisdom R, Johnson RS, Moore C. c-Jun regulates cell cycle progression and apoptosis by distinct mechanisms. EMBO J 1999;18:188–97.

[48] Taylor-Harding B, Aspuria PJ, Agadjanian H, Cheon DJ, Mizuno T, Greenberg D, et al. Cyclin E1 and RTK/RAS signaling drive CDK inhibitor resistance via activation of E2F and ETS. Oncotarget 2015;6:696–714.

[49] Wang B, Krall EB, Aguirre AJ, Kim M, Widlund HR, Doshi MB, et al. ATXN1L, CIC, and ETS Transcription Factors Modulate Sensitivity to MAPK Pathway Inhibition. Cell Reports 2017;18:1543–57.

[50] Tadesse S, Yu M, Kumarasiri M, Le BT, Wang S. Targeting CDK6 in cancer: State of the art and new insights. Cell Cycle 2015;14:3220–30.

[51] Hugo W, Shi H, Sun L, Piva M, Song C, Kong X, et al. Non-genomic and Immune Evolution of Melanoma Acquiring MAPKi Resistance. Cell 2015;162:1271–85.

[52] Gowrishankar K, Snoyman S, Pupo GM, Becker TM, Kefford RF, Rizos H. Acquired Resistance to BRAF Inhibition Can Confer Cross-Resistance to Combined BRAF/MEK Inhibition (vol 132, pg 1850, 2012). Journal of Investigative Dermatology 2013;133:2493-.

[53] Hou P, Wu C, Wang Y, Qi R, Bhavanasi D, Zuo Z, et al. A Genome-Wide CRISPR Screen Identifies Genes Critical for Resistance to FLT3 Inhibitor AC220. Cancer Res 2017;77:4402–13.

[54] Shalem O, Sanjana NE, Hartenian E, Shi X, Scott DA, Mikkelsen TS, et al. Genome-Scale CRISPR-Cas9 Knockout Screening in Human Cells. Science 2014;343:84–7.

[55] Singleton KR, Crawford L, Tsui E, Manchester HE, Maertens O, Liu X, et al. Melanoma Therapeutic Strategies that Select against Resistance by Exploiting MYC-Driven Evolutionary Convergence. Cell Rep 2017;21:2796–812.

[56] Asghar U, Witkiewicz AK, Turner NC, Knudsen ES. The history and future of targeting cyclin-dependent kinases in cancer therapy. Nat Rev Drug Discov 2015;14:130–46.

[57] Sondergaard JN, Nazarian R, Wang Q, Guo D, Hsueh T, Mok S, et al. Differential sensitivity of melanoma cell lines with BRAFV600E mutation to the specific Raf inhibitor PLX4032. J Transl Med 2010;8:39.

[58] Sanjana NE, Shalem O, Zhang F. Improved vectors and genome-wide libraries for CRISPR screening. Nat Methods 2014;11:783–4.

[59] Ritchie ME, Phipson B, Wu D, Hu Y, Law CW, Shi W, et al. limma powers differential expression analyses for RNA-sequencing and microarray studies. Nucleic Acids Res 2015;43:e47.

[60] Corces MR, Trevino AE, Hamilton EG, Greenside PG, Sinnott-Armstrong NA, Vesuna S, et al. An improved ATAC-seq protocol reduces background and enables interrogation of frozen tissues. Nat Methods 2017;14:959–62.

[61] Qin Q, Mei S, Wu Q, Sun H, Li L, Taing L, et al. ChiLin: a comprehensive ChIP-seq and DNase-seq quality control and analysis pipeline. BMC Bioinformatics 2016;17:404.

[62] Quinlan AR, Hall IM. BEDTools: a flexible suite of utilities for comparing genomic features. Bioinformatics 2010;26:841–2.

[63] Love MI, Huber W, Anders S. Moderated estimation of fold change and dispersion for RNA-seq data with DESeq2. Genome Biol 2014;15:550.

[64] Wang S, Sun H, Ma J, Zang C, Wang C, Wang J, et al. Target analysis by integration of transcriptome and ChIP-seq data with BETA. Nat Protoc 2013;8:2502–15.

[65] Ianevski A, He L, Aittokallio T, Tang J. SynergyFinder: a web application for analyzing drug combination dose-response matrix data. Bioinformatics 2017;33:2413–5.

[66] Warde-Farley D, Donaldson SL, Comes O, Zuberi K, Badrawi R, Chao P, et al. The GeneMANIA prediction server: biological network integration for gene prioritization and predicting gene function. Nucleic Acids Res 2010;38:W214–20.

